# A bias in transsaccadic perception of spatial frequency changes

**DOI:** 10.1101/2024.03.11.584439

**Authors:** Nino Sharvashidze, Carolin Hübner, Alexander C. Schütz

**Author notes:** Corresponding author. E-mail address, Sensomotoric Learning, Experimental & Biological Psychology, Department of Psychology, Philipps-Universität Marburg, Gutenbergstraße 18, 35032 Marburg, Germany.

## Abstract

Visual processing differs between the foveal and the peripheral visual field. These differences can lead to different appearances of objects in the periphery and the fovea, which poses a challenge to perception across saccades. Differences in the appearance of visual features between the peripheral and foveal visual field may bias change discrimination across saccades. Previously it has been reported that spatial frequency (SF) appears higher in the periphery compared to the fovea (Davis et al., 1987). In this study, we investigated the visual appearance of SF before and after a saccade and the discrimination of SF changes implemented during saccades. In addition, we tested the contributions of pre- and postsaccadic information to change discrimination performance. In the first experiment, we found no differences in the appearance of SF before and after a saccade. However, participants showed a clear bias to report SF increases. Interestingly, a 200-ms postsaccadic blank period improved the precision of the responses but did not affect the bias. In the second experiment, participants showed lower thresholds for SF increases than for decreases, suggesting that the bias in the first experiment was not just a response bias. Finally, we asked participants to discriminate the SF of stimuli presented before a saccade. Thresholds in the presaccadic discrimination task were lower than thresholds in the change discrimination task, suggesting that transsaccadic change discrimination is not merely limited by presaccadic discrimination in the periphery. The change direction bias might stem from more effective masking or overwriting of the presaccadic stimulus by the postsaccadic low SF stimulus.

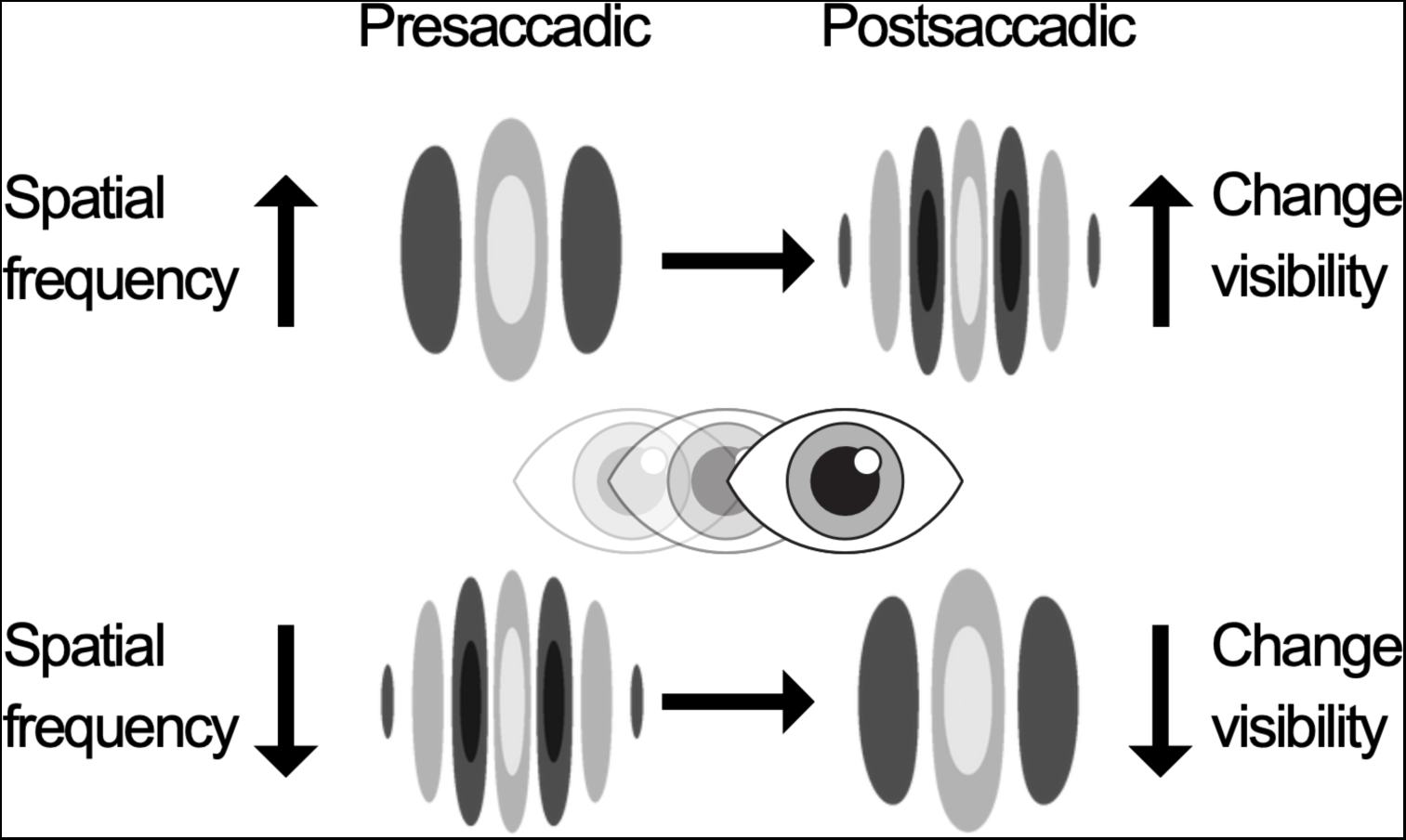

## 1. Introduction

High-resolution vision is primarily located in the fovea due to its dense photoreceptor distribution. Peripheral vision provides a broader field of view, at the cost of lower resolution and contrast sensitivity, and increased spatial distortion (for reviews see Rosenholtz, 2016; Strasburger et al., 2011). Various mechanisms operate to align the appearance of the fovea and the periphery, resulting in a rather homogeneous perception across the visual field (for reviews see Cohen et al., 2016; Knotts et al., 2019; Stewart et al., 2020). Despite these mechanisms, the appearance of visual features of objects is not perfectly matched between the periphery and the fovea. For example, object size (Newsome, 1972) and shape (Baldwin et al., 2016; Coates et al., 2017; Hübner & Schütz, 2021; Valsecchi et al., 2018) have been reported to appear differently in the periphery and at the fovea. Spatial frequency (SF) has been reported to appear higher in the periphery compared to the fovea (Davis et al., 1987; Georgeson, 1980; Harris & Wink, 2000). In foveal vision, sensitivity is highest for intermediate SFs (approximately 2 to 8 cycles per degree [cpd]) and is reduced for lower and higher SFs (Contrast Sensitivity Function [CSF]; Campbell & Robson, 1968). Due to spatial pooling in peripheral vision, the CSF retains a similar shape but is shifted towards lower SFs (Rovamo et al., 1978). Therefore, the perception of low SFs with increasing eccentricity is largely unaffected, while detecting high SFs becomes challenging. The overestimation of SF in the periphery has been attributed to the possibility that the labels for SF channels do not update to compensate for this shift (Davis, 1990; Georgeson, 1980).

The differences between visual processing and appearance across the visual field lead to challenges for perception across saccades. Reduced processing quality in the periphery requires recurrent saccadic eye movements to take advantage of foveal acuity at different locations in the environment (e.g. Rayner, 1998). The visual system faces the task of tracking, comparing, and matching object identity, location, and features across every eye movement. If the differences in the processing or appearance of visual features are inherently consistent, then specific feature changes, and even specific directions of feature change, are more or less likely to occur with an eye movement. It has been shown that these regularities do not go unnoticed. Through transsaccadic learning, contingencies are learned and predictions about the saccade outcome are formed (Herwig & Schneider, 2014). Experiments systematically manipulating pre- and postsaccadic contingencies have shown that transsaccadic predictions influence presaccadic perception and probably play a role in the recalibration of peripheral and foveal appearance (e.g. Bosco et al., 2015; Cox et al., 2005; Herwig & Schneider, 2014; Herwig et al., 2018; Hübner & Schütz, 2021; Köller et al., 2020; Paeye et al., 2018; Valsecchi & Gegenfurtner, 2016). In addition to the recalibration function, transsaccadic prediction could directly impact the perception of visual features during saccades. A viable method to investigate this influence involves inducing changes in specific features during saccades and subsequently assessing the capacity to discriminate these changes (Hübner & Schütz, 2021). For instance, if the SF of an object appears to be higher before a saccade than after, it may create a general prediction that SF decreases after a saccade. It could be that perception along the prediction, i.e. SF decrease perception, is facilitated. It is also plausible that a change contradicting the prediction enhances change perception through prediction error, making an increase in SF more readily detectable.

Hübner and Schütz (2021) found a relationship between the inherent appearance differences of participants and their bias in detecting changes across saccades. Observers on average perceived presented object shapes as more circular in presaccadic, peripheral view compared to postsaccadic foveal view. Across saccades, physical shape changes opposite to this regularity, namely circularity-increasing changes, were more likely to be detected.

In the present study, we investigated the perception of spatial-frequency changes during saccades. Building on previous research showing overestimation of SF in peripheral vision (Davis et al., 1987; Georgeson, 1980; Harris & Wink, 2000), our primary aim was to investigate whether SF appearance differs before and after saccades and whether this discrepancy affects transsaccadic change discrimination, as shown for shape by Hübner and Schütz (2021). We additionally incorporated a postsaccadic blank in the change discrimination task. Temporarily removing the saccade target during eye movement has been shown to enhance transsaccadic change detection (e.g. Deubel et al., 1996; Weiss et al., 2015).

In the first experiment, we investigated the appearance of SF before and after a saccade and compared the discrimination of changes in two directions: SF increase and decrease. We did not find any consistent differences in apparent SF before and after a saccade. However, we observed a bias to detect an increase in SF. Introducing a 200-ms postsaccadic blank improved the change discrimination but did not alleviate the bias. In our second experiment, we aimed to better understand the source of the bias as it remained unexplained by appearance differences. Here, we accounted for a potential response bias and included a presaccadic SF discrimination task. This enabled us to compare discrimination thresholds before and across a saccade. The lower thresholds in the presaccadic SF discrimination task suggest that perception is hindered at the time when the postsaccadic input arrives. This hindrance might come from masking/overwriting of the presaccadic information.

## 2. Experiment 1

### 2.1 Methods

#### 2.1.1 Participants

In total, 18 participants took part in Experiment 1. One participant had to be excluded due to a misunderstanding of the task instructions. The data of 17 participants (3 males, mean age = 23.94 ± 5.13 years, range 19-42 years) were used for the analysis. All participants were unaware of the purpose of the study and were paid 8€ per hour. They had normal or corrected-to-normal vision and gave informed consent before participating. The experiment was conducted in accordance with the principles of the 1964 Declaration of Helsinki and was approved by the ethics committee of the Philipps University of Marburg, Department of Psychology (proposal 2015-35k).

#### 2.1.2 Stimuli

The fixation stimulus was a bull’s-eye and crosshair combination (Thaler et al., 2013), spanning 0.6 degrees of visual angle (dva). The color of the fixation stimulus was randomly selected from a DKL color-space array (Derrington et al., 1984), varying in the polarity of the luminance and red-green color channel to avoid the build-up of after images. As discrimination stimuli, we used Gabor patches with randomized orientations (−180° to 180°) and phases, surrounded by a Gaussian envelope with a standard deviation of 0.5 dva. All stimuli were shown on a gray (0.5) background. The Gabors had 50% contrast and were presented at ±15 dva eccentricity on the horizontal axis from the central fixation stimulus. The SF of the Gabors was adjusted in quarter-octave increments around 2.0 cpd, resulting in the subsequent SF spectrum: 0.84, 1.0, 1.19, 1.41, 1.68, 2.0, 2.38, 2.83, 3.36, 4.0, 4.76 cpd. A black dot of 0.15 dva diameter was located at the center of the Gabor.

#### 2.1.3 Design

In the appearance discrimination task, the Gabor stimuli were presented either only before saccades, followed by a postsaccadic fixation cross, or only after saccades, following a presaccadic fixation cross. Participants were asked to determine whether the SF in each trial was higher or lower than the mean SF of all stimuli observed until that trial, utilizing the method of single stimuli as in previous studies (Hübner & Schütz, 2017; Hübner & Schütz, 2021; Morgan et al., 2000). For both the presaccadic and postsaccadic conditions, eleven SF values were examined, each repeated 15 times, resulting in a total of 330 trials.

In the change discrimination task of the first experiment, the SF could either increase or decrease during a saccade. Participants were asked to identify the direction of SF change. In half of the trials, a 200 ms blank screen was presented with saccade detection. For each change direction and blanking condition, two separate staircases were employed. One of these staircases was initiated with the smallest change magnitude of 0.25 |Δ octave units|, while the other with the largest magnitude of 2.50 |Δ octave units|. The presaccadic SF was randomly selected from a range of spatial frequencies that were suitably distant from the upper and lower limits of the frequency range, considering the desired change magnitude and direction. For example, if the change magnitude of a trial was 2.0 |Δ octave units| and the direction of change was SF increase, the presaccadic value was randomly selected from 0.84, 1.0 and 1.19 cpd. Responses with the reported change not matching the actual change direction were classified as a miss, resulting in an increase of 0.25 |Δ octave units| in magnitude for the subsequent trial. Responses were classified as a hit if the reported and actual change directions matched. After achieving two consecutive hits, the change magnitude was decreased. Each staircase was repeated 50 times, resulting in a total of 400 trials. A training session was introduced with a design identical to the main experiment. It consisted of 4 staircase repetitions, resulting in 32 training trials. Additionally, a high-pitched tone feedback was incorporated into the training session as a response to incorrect answers. Training trials were not included in the analysis. Trials were randomly interleaved across all conditions in both tasks. The order in which the tasks were conducted was counterbalanced among participants.

#### 2.1.4 Equipment

Stimuli were presented on a VIEWPixx monitor at a viewing distance of 60 cm. The size of the monitor was 51.5 x 29 cm, with a resolution of 1920 × 1080 pixels and a refresh rate of 120 Hz (VPixx Technologies Inc., Quebec, Canada). The screen was calibrated to achieve a linear gamma correction (luminance values 0.39, 54, 105 cd/m² for black, gray, and white). Eye movements were recorded using a desktop-mounted EyeLink 1000 Plus system (SR Research Ltd., Ontario, Canada) at 1000 Hz. MATLAB (R2017a) with Psychophysics Toolbox (3.0.12) (Brainard, 1997; Pelli, 1997) was used for the stimulus display, and the EyeLink Toolbox (Cornelissen et al., 2002) was used for the eye tracker operation. Head movement was restricted by a chinrest. Participants responded using a standard keyboard (vertical plus-sign key on number pad for frequency higher than mean/frequency decrease and horizontal zero key on number pad for lower than mean/ frequency decrease responses). Data analysis was conducted with MATLAB (R2021a) and RStudio (4.3.0).

#### 2.1.5 Procedure

Participants initiated each trial by pressing the space bar while fixating on the central fixation stimulus. In the appearance discrimination task, the presaccadic stimulus could either be a fixation cross (in the postsaccadic condition) or a Gabor (in the presaccadic condition) (Fig. 1A). The presaccadic stimulus appeared to the left or right on the horizontal axis at an eccentricity of 15 dva after a variable period between 750 and 1500 ms. Participants were instructed to make a saccade to the stimulus. Upon saccade detection, the presaccadic stimulus either changed into a Gabor (in the postsaccadic condition) or a fixation stimulus (in the presaccadic condition). The fixation stimulus at the center remained on the screen for an additional 200 ms after target onset or until a saccade was detected. Saccades were continuously monitored online and their onset was determined as the moment when the gaze position exceeded a circular area of 2.0 dva radius centered around the central fixation stimulus. A trial was aborted when no saccade was detected within 1.8 seconds after saccade target onset. Saccades landing outside a 2 dva region of the center of the Gabor and latency errors (saccade latencies falling below 50 ms or exceeding 600 ms) were followed by a low-pitched tone. Postsaccadic stimulus presentation duration was half the participant’s median presaccadic stimulus presentation duration over all presaccadic conditions of the completed trials. After the disappearance of the postsaccadic stimulus, participants were required to indicate whether the SF of the Gabor in that trial was higher (vertical plus-sign key) or lower (horizontal zero key) compared to the mean SF of all stimuli seen until that point. There was no blank condition in the appearance discrimination task.

**Fig. 1.**
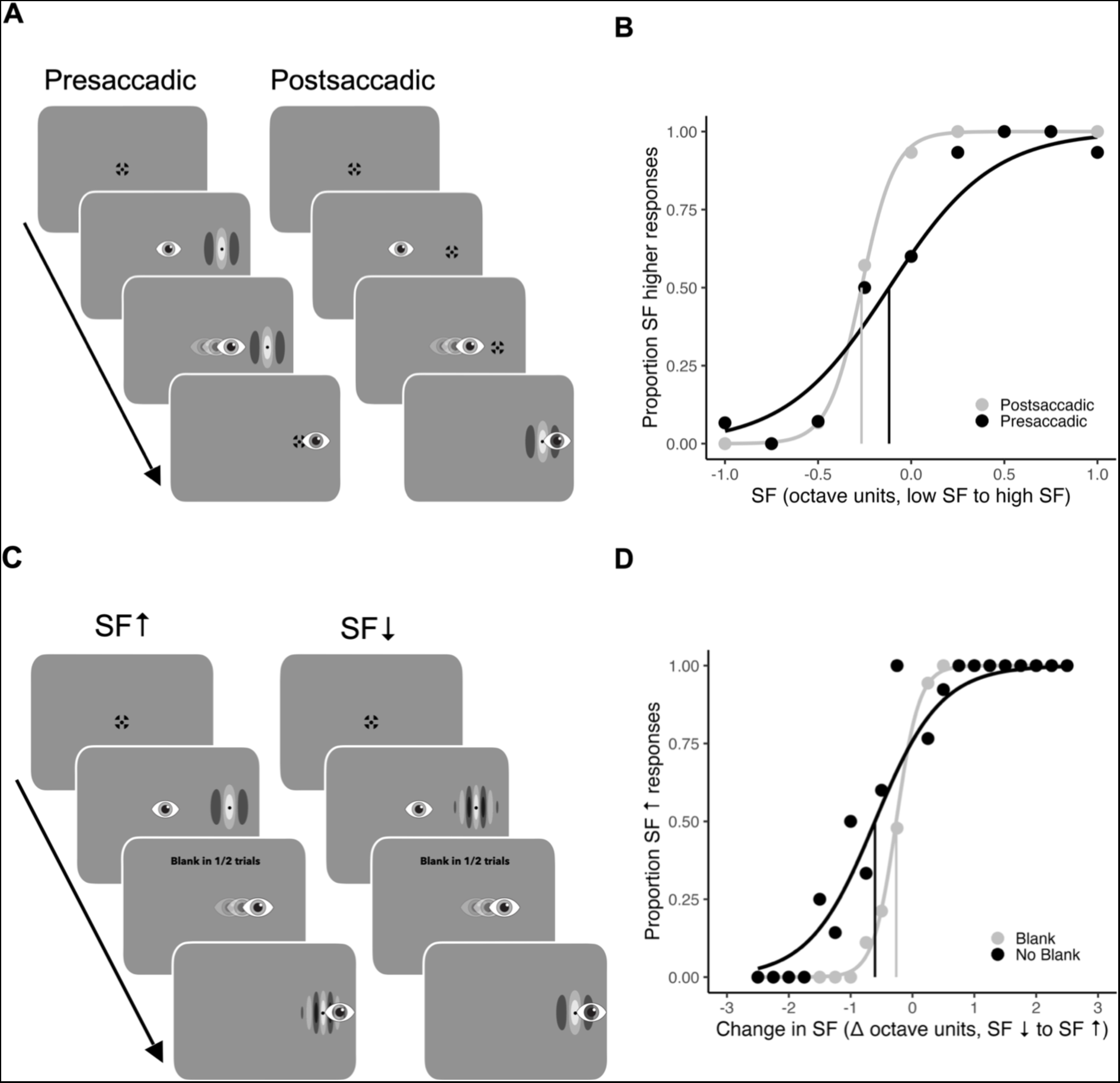
Stimuli and methods of Experiment 1. A) Schematic trial procedure of Experiment 1, Appearance discrimination task. Participants had to compare the SF of the Gabor in this trial to the overall mean SF across all previous trials. The Gabors were either presented before the saccade in the peripheral visual field (presaccadic condition) or after the saccade in the central visual field (postsaccadic condition). B) Example psychometric functions of one representative participant fitted to proportion SF-higher responses over SF in octave units for presaccadic (black) and postsaccadic (gray) conditions. Zero indicates the mean SF in octave units (2 cpd), negative values indicate lower SFs and positive values indicate higher SFs. Vertical lines indicate the points of subjective equality (PSE). C) Schematic trial procedure of Experiment 1, Change discrimination task. The SF of the Gabor either increased or decreased during the saccade and participants had to indicate the direction of change. D) Example psychometric functions of one representative participant fitted to proportion SF-increase responses over SF changes tested for blank (gray) and no blank (black) conditions. Negative deltas indicate SF decrease and positive deltas indicate SF increase. Vertical lines indicate points of subjective stability (PSS).

The change discrimination task had a similar trial procedure except that the target stimulus was always a Gabor (Fig. 1C). Upon saccade detection, the presaccadic Gabor was replaced either immediately (no blank condition), or removed for 200 ms (blank condition) and then replaced by the postsaccadic Gabor. The duration of the postsaccadic stimulus display equaled half that of the presaccadic stimulus for a given trial. After the disappearance of the postsaccadic stimulus, participants were required to indicate whether the SF of the target had increased (vertical plus-sign key) or decreased (horizontal zero key) during their saccade.

#### 2.1.6 Eye movement analysis and trial exclusions

For the analysis of eye movement data, saccades were identified offline using the EyeLink algorithm (velocity threshold = 22°/s, acceleration threshold = 3800°/s²). Saccade landing position was the gaze position at saccade offset. Saccade latency was the time between presaccadic stimulus onset and saccade onset. Trials were excluded if participants blinked during the period from 300 ms before target onset until target offset, if the pre- to postsaccadic stimulus switch did not happen during the saccade, or if the full sequence of events was not completed (0.45 ± 0.94% of trials in the appearance discrimination task and 0.28 ± 0.52% in the change discrimination task). Additionally, trials with saccade latencies below 50 ms or above 600 ms (4.05 ± 4.73% in the appearance discrimination task and 4.43 ± 4.41% in the change discrimination task) and those where the gaze position deviated more than 2 dva on the horizontal axis or 1.5 dva on the vertical axis from the saccade target center between saccade offset and target offset were also excluded (9.20 ± 5.70% in the appearance discrimination task and 7.90 ± 5.96% in the change discrimination task). In total, 10.23 ± 6.3% of trials were excluded from the appearance discrimination task and 9.97 ± 6.57% from the change discrimination task.

#### 2.1.7 Psychophysical analysis

To derive psychometric functions in the appearance discrimination task, the responses were transformed into proportions of responses indicating higher SF compared to the mean, for each SF that was tested. Psychometric functions were then fitted, with SF in octave units (Fig. 1B). The point of subjective equality (PSE) was estimated as the SF corresponding to 50% SF higher (than mean) responses. A PSE above zero indicates a tendency to perceive SF as lower, while a PSE below zero indicates a tendency toward perceiving it as higher. The just-noticeable difference (JND) was defined as half of the difference between the thresholds at 25% and 75%, with a lower JND denoting greater precision in discriminating higher/lower SF. In the change discrimination task, we converted each participant’s perceptual choices into proportion SF-increase responses for each tested change measured by the difference between octave units (Fig. 1D). The point of subjective stability (PSS) was estimated as the magnitude and direction of SF change corresponding to 50% SF-increase responses. A negative PSS indicates a bias toward reporting SF increases. The JND serves as an indicator of an observer’s ability to detect changes and was calculated as half of the difference between the thresholds at 25% and 75% perception levels. A lower JND suggests greater precision in discriminating SF changes. We employed the Quickpsy package in R (Linares & López-Moliner, 2016) to fit a logistic function to the data of both tasks. We report t-tests, p values, and Bayesian t-tests (BF_10_ indicates evidence against the null hypothesis). T-tests were calculated in R using the stats package (R Core Team, 2023). Two-sided t-tests were conducted at a significance level of 0.05. Bayesian t-tests were performed using the BayesFactor package in R (Morey & Rouder, 2022).

### 2.2 Results

In the appearance discrimination task, we measured the appearance of SF before and after a saccade. The mean point of subjective equality (PSE) was −0.18 ± 0.23 octave units in the presaccadic condition and −0.21 ± 0.17 octave units in the postsaccadic condition. The mean PSEs in the two conditions did not differ from each other (t(16) = 0.71, p = 0.485, BF_10_ = 0.31, anecdotal evidence for H0) (Fig. 2A). The mean just-noticeable difference (JND) in the presaccadic condition was 0.34 ± 0.17 octave units and 0.23 ± 0.12 octave units in the postsaccadic condition. There was a significant difference between pre- and postsaccadic conditions (t(16) = 3.94, p = 0.001, BF_10_ = 32.72, very strong evidence for H1) indicating higher precision in the postsaccadic condition (Fig. 2B).

**Fig. 2.**
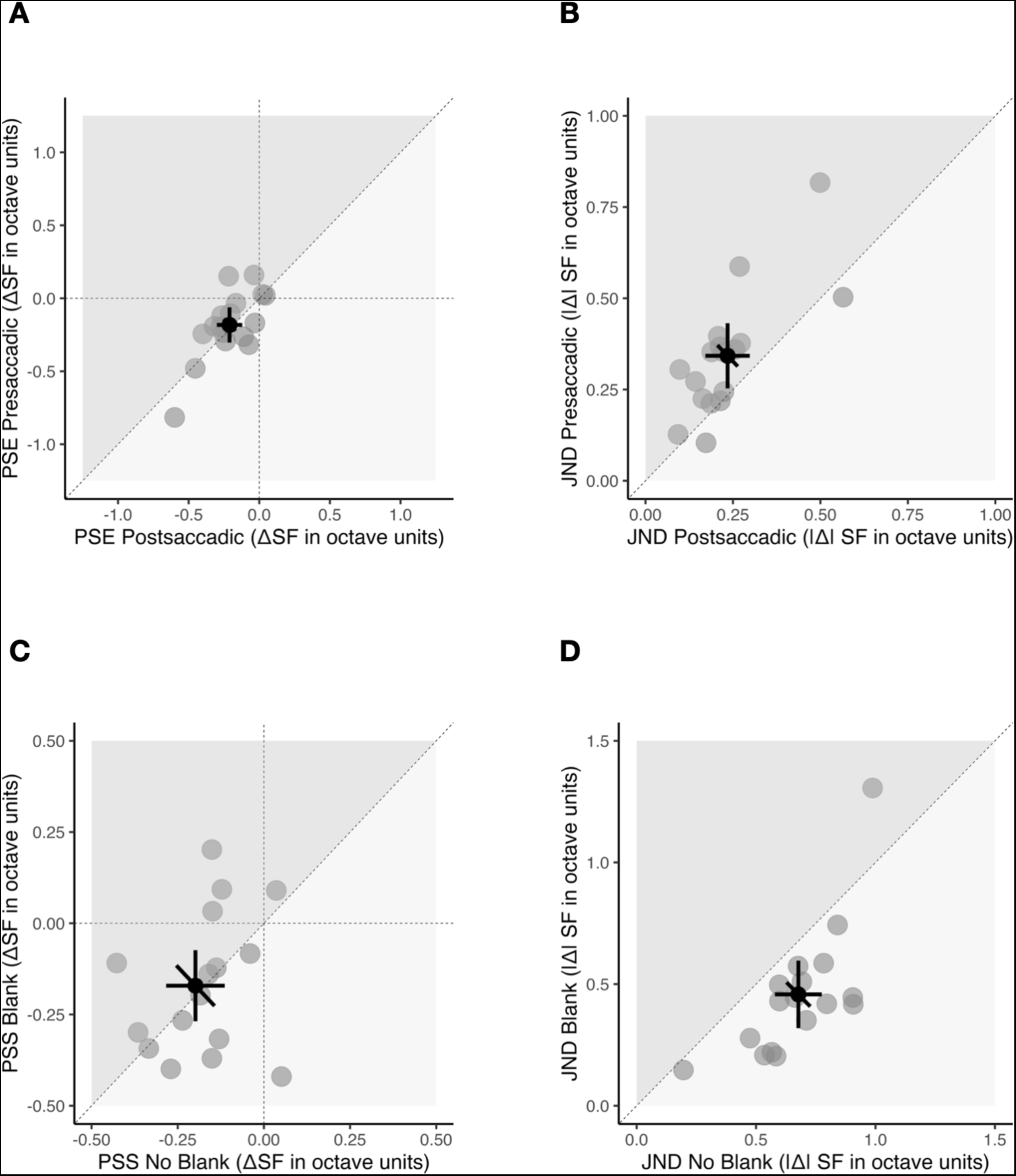
Results of Experiment 1. A & B) Appearance discrimination task. C & D) Change discrimination task. A) Scatterplot for all points of subjective equality (PSE) compared between presaccadic (vertical axis) and postsaccadic (horizontal axis) conditions. Data points on the dashed diagonal line indicate no difference in SF appearance between the presaccadic and postsaccadic conditions. B) Scatterplot for all just-noticeable differences (JNDs) compared between presaccadic (vertical axis) and postsaccadic (horizontal axis) conditions. Data points above the diagonal dashed line indicate that participants were less precise in the presaccadic condition. C) Scatterplot for all points of subjective stability (PSS) compared between the blank (vertical axis) and no blank (horizontal axis) conditions. Data points left from the dashed vertical line or below the dashed horizontal line (negative PSS) indicate a bias for SF-increase changes. D) Scatterplot for all just-noticeable differences (JNDs) compared between blank (vertical axis) and no blank (horizontal axis) conditions. Data points below the diagonal dashed line indicate that participants were less precise in the no blank condition. A - D) The individual participant data are represented by light gray dots and the overall mean is indicated by a dark gray dot. The error bars show 95%-confidence intervals for each condition (cardinal bars) or between conditions (oblique bar).

In the change discrimination task, we increased or decreased SF of the Gabor during the saccade and asked participants to report the perceived direction of the change. The mean point of subjective stability (PSS) in the blank condition was −0.17 ± 0.19 ΔSF octave units and −0.2 ± 0.17 ΔSF octave units in the no blank condition. The negative values indicate a bias to report SF-increase changes in both no blank and blank conditions (Fig. 2C). The mean PSS in the no blank condition significantly deviated from zero (t(16) = −4.96, p < 0.001, BF_10_ = 208.09, extreme evidence for H1). Similarly, the PSS in the blank condition was significantly different from zero (t(16) = −3.73, p = 0.002, BF_10_ = 22.52, strong evidence for H1). There was no difference between the PSS in the blank and no blank conditions (t(16) = 0.53, p = 0.61, BF_10_ = 0.36, anecdotal evidence for H0). The mean JND for SF change discrimination in the no blank condition was 0.68 ± 0.19 |ΔSF octave units| and 0.46 ± 0.27 |ΔSF octave units| for the blank condition. JNDs were significantly different (t(16) = −4.70, p = 0.0003, BF_10_ = 129.74, extreme evidence for H1) between the two blanking conditions (Fig. 2D). In sum, participants were significantly more precise (JNDs) but not more accurate (PSS) at discriminating SF changes in the blank condition.

## 3. Experiment 2

In the first experiment, no differences were observed in SF appearance before and after a saccade. Moreover, there was no correlation between the appearance and participants’ change discrimination performance (see Supplementary Material). Nevertheless, participants showed a bias to report an increase in SF across saccades. Firstly, it is not entirely clear whether this observed directional bias is indeed a perceptual bias or whether it can be attributed to a preference for a particular change direction response or key press. To address this question, we conducted a second experiment with a change discrimination task that did not rely on predefined criteria for change direction. Participants were asked to report the location rather than the direction of the change. In addition, we wanted to ensure that the bias did not come from the poor presaccadic SF discrimination, as sensitivity for high SF is reduced in the periphery (Rovamo et al., 1978). To this end, we introduced a task where participants had to discriminate SF based on presaccadic information only. As a result, we could additionally compare thresholds between presaccadic SF discrimination (Experiment 2, presaccadic discrimination task) and transsaccadic SF change (Experiment 2, change discrimination task) conditions. If the bias in the case of SF cannot be adequately explained by predictions based on appearance differences, there may be alternative mechanisms at play. Examining the separate influences of the two information sources can provide insight into the underlying mechanisms involved.

To understand the limitations and biases of change perception, it is essential to understand how information is processed across saccades. Presaccadic information about SF (Weiss et al., 2015), orientation (Ganmor et al., 2015; Grzeczkowski, Deubel, & Szinte, 2020; Stewart & Schütz, 2018; Wolf & Schütz, 2015), and other visual features of the saccade target can be transferred across saccades and integrated with postsaccadic information (shape - Demeyer et al., 2011; Grzeczkowski, van Leeuwen, et al., 2020; numerosity - Hübner & Schütz, 2017; color - Schut et al., 2018; Wijdenes et al., 2015; Wittenberg et al., 2008). At the same time, the precise nature of transsaccadic integration remains uncertain. Some findings suggest a basis in abstract visual memory (Prime et al., 2006; Stewart & Schütz, 2018), while others argue for reliance on highly detailed sensory representation. For instance, certain conditions allow for the system to generate a low-level pattern overlay through the fusion of pre- and postsaccadic information (Paeye et al., 2017). This information transfer is primarily low-level, suggesting it can be susceptible to masking or overwriting.

Masking/overwriting has been reported with different tasks. Transsaccadic spatiotopic backward masking has been reported in a perisaccadic target detection task (De Pisapia et al., 2010). Furthermore, in transsaccadic tasks where retaining separate pre- and postsaccadic representations is crucial, presaccadic information can be overwritten by postsaccadic information (Tas et al., 2012; Tas et al., 2021). It has been proposed that object continuity is responsible for the overwriting mechanism whereby the postsaccadic value of the target overwrites the presaccadic value, provided both are perceived as properties of a single, continuous object (Tas et al., 2012). A similar explanation comes from the theory of task-driven visual attention and working memory (TRAM), which highlights the importance of assessing object continuity and correspondence in determining attentional priorities during saccades (Poth & Schneider, 2016; Schneider, 2013). The concept of object continuity also aligns with the blanking phenomenon (Deubel et al., 1996). The presence of a blank period potentially facilitates the retrieval of information present before the saccade that is otherwise overwritten by or integrated with the postsaccadic information (De Pisapia et al., 2010; Deubel et al., 1996; Tas et al., 2021). Another line of research has shown that perception of shifts in object position is also impaired during eye movements resulting in difficulty to accurately identify displacement directions (saccadic suppression of displacement [SSD]; Bridgeman et al., 1975; Bridgeman & Stark, 1979). Masking has been recently posited as a potential cause of the SSD (Takano et al., 2020). Takano et al. (2020) showed that introducing a high-contrast stimulus after a saccade increased SSD. Accessing the separate influences of the pre- and postsaccadic inputs on change discrimination in the second experiment allows to examine the potential role of masking/overwriting in SF change discrimination.

### 3.1 Methods

#### 3.1.1 Participants

In Experiment 2, a group of thirteen participants took part, of whom two had previously participated in Experiment 1. One participant did not complete both tasks. We had to exclude four further participants from the analysis. These participants were unable to achieve 75% correct responses in one or more conditions within one or both tasks, even with the largest SF changes/differences. The data of eight participants (two males, mean age = 25 ± 4.96 years, range 17-32 years) was used for the analysis. All participants were unaware of the purpose of the studies and were compensated with 8€ per hour.

#### 3.1.2 Stimuli

In Experiment 2, two Gabors were presented simultaneously, one below and one above a fixation stimulus centered between them, with 2.5 dva between the center of one Gabor and the center of the fixation stimulus. Due to the challenging nature of this change discrimination task, we modified the eccentricity, contrast settings, and SF range compared to the first experiment. The Gabors had 80% contrast and were shown at ± 5 dva eccentricity on the horizontal axis from the central fixation stimulus. The first task employed one-third-octave increments around 2 cpd resulting in the following values: 0.71, 0.87, 1.07, 1.32, 1.62, 2.0, 2.46, 3.03, 3.73, 4.59, 5.66 cpd. The presaccadic discrimination task was less challenging and employed 0.05-octave increments around 2 cpd resulting in the following values: 1.68, 1.74, 1.80, 1.87, 1.93, 2.0, 2.07, 2.14, 2.23, 2.30, 2.38 cpd. The contrast and eccentricity had to remain the same.

#### 3.1.3 Design

The change discrimination task of Experiment 2 was similar to the change discrimination task of Experiment 1, except here two Gabors with different SFs were presented before a saccade and the SF of one of them changed during a saccade, resulting in both Gabors having the same SF after a saccade. Participants were instructed to indicate whether the SF of the top- or the bottom Gabor was changed. Four staircases were repeated 70 times, resulting in 560 trials. A high-pitched tone signaled incorrect responses, while a low-pitched tone indicated a saccade landing outside a 2 dva region from the fixation cross center. Feedback aimed to improve participant performance and prevent saccades from landing on the Gabor gratings. In the second, presaccadic discrimination task, two Gabors of differing SFs and a fixation cross were shown before a saccade and the Gabors disappeared upon the detection of a saccade. The participants’ task was to identify whether the upper or the lower Gabor had a higher SF. Two staircases were repeated 80 times resulting in 320 trials. A high-pitched tone signaled incorrect responses. In the change discrimination task, a training session was introduced with a design identical to the main experiment. It consisted of four staircase repetitions, resulting in 32 training trials. Training trials were not included in the analysis. Trials in both tasks were randomly interleaved across all conditions. The order in which the tasks were conducted was counterbalanced across participants.

#### 3.1.4 Equipment

The equipment was identical to that described in Experiment 1. Participants responded using up- and down arrow keys for top/bottom responses.

#### 3.1.5 Procedure

During the change discrimination task of the second experiment, participants fixated on the central fixation stimulus for a period ranging from 750 to 1500 ms. Following this, two presaccadic Gabors and a fixation stimulus appeared to the left or right on the horizontal axis at an eccentricity of 5 dva. Participants were instructed to make a saccade towards the fixation stimulus between the Gabors. Upon detecting a saccade, one of the Gabors underwent a SF change, aligning itself with the second Gabor after the saccade. The change occurred either immediately or after a 200 ms blank interval. The duration of the postsaccadic stimulus was half of the presaccadic stimulus duration for each trial. After the stimuli disappeared, participants were asked to indicate which of the two Gabors (upper or lower) had undergone the SF change by using the up and down arrow keys (Fig. 3A). The presaccadic discrimination task followed a similar trial procedure, with the exception that Gabors disappeared upon saccade detection, leaving only the fixation stimulus on the screen. Participants were then asked to indicate which of the two presaccadic Gabors had a higher SF by using the upper and lower arrow keys (Fig. 3C).

**Fig. 3.**
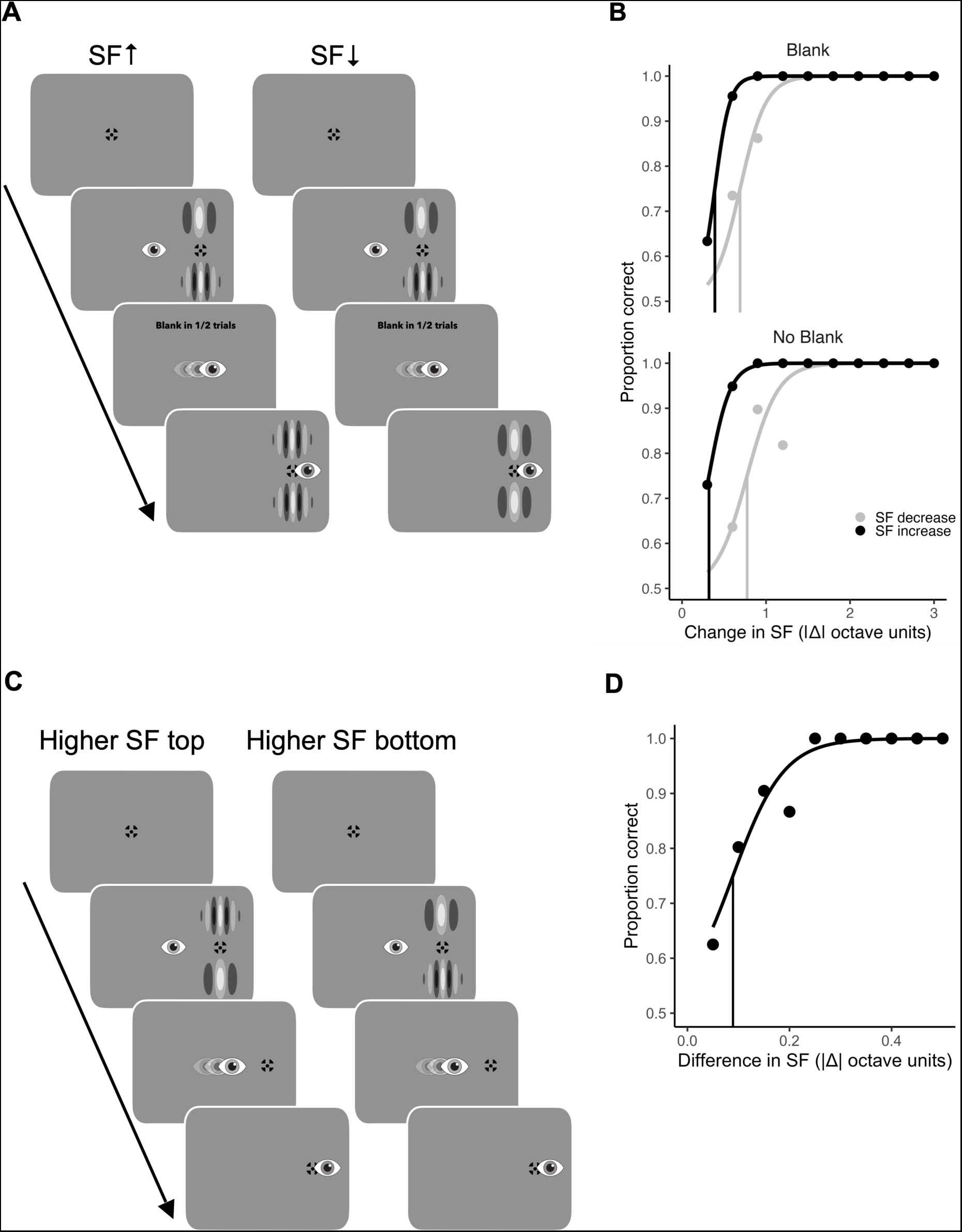
Stimuli and methods of Experiment 2. A) Schematic trial procedure of Experiment 2, Change discrimination task. Two SF Gabors were presented presaccadically and one of them changed its SF during the saccade, resulting in both Gabors showing the same SF after a saccade. Participants had to indicate whether the change happened at the top or at the bottom Gabor. B) Example psychometric functions of one representative participant fitted to proportion correct responses over absolute SF change magnitudes in octave units for SF increase (black) and decrease (gray) conditions. Blank condition at the top, no blank condition at the bottom. Vertical lines indicate thresholds at 75% correct responses. C) Schematic trial procedure of Experiment 2, Presaccadic discrimination task. Two SF Gabors were presented presaccadically and disappeared after a saccade. Participants had to indicate whether the higher SF was at the top or the bottom. D) Example psychometric function of one representative participant fitted to proportion correct responses over the absolute difference in SF in octave units. The vertical line indicates the threshold at 75% correct responses.

#### 3.1.6 Eye movement analysis and trial exclusions

The analysis and trial exclusion criteria were identical to Experiment 1. In total, 8.84 ± 6.33% of the trials were excluded from the first task and 14.34 ± 11.29% from the second task.

#### 3.1.7 Psychophysical analysis

Participants’ perceptual choices were transformed into proportions of correct responses for various magnitudes of SF changes (difference between octave units) in both change-direction conditions in the change discrimination task (Fig. 3B) and for the difference in SF between the presaccadic stimuli in the presaccadic discrimination task (Fig. 3D). A logistic function was applied to the data for each participant, beginning at a baseline chance level of 50% correct responses. The threshold for detection was calculated as the absolute SF change magnitude required for a participant to achieve a 75%-correct response rate. A lower threshold value indicates greater sensitivity to the corresponding SF-change direction.

### 3.2 Results

To show that the change-direction bias in the first experiment was not a mere bias toward a particular response option or key, we conducted Experiment 2. In the first task, participants were presented with a pair of Gabor stimuli before and after a saccade, with only one of them changing its SF during the saccade. The mean change detection threshold across both change direction conditions was 0.79 ± 0.23 |ΔSF octave units|. In the SF decrease condition, the mean change detection threshold was 1.00 ± 0.27 |ΔSF octave units| and in the SF increase condition it was 0.57 ± 0.18 |ΔSF octave units|. Detection thresholds were significantly lower in the SF increase condition (t(7) = 7.72, p < 0.001, BF_10_ = 245.23, extreme evidence for H1) (Fig. 4A). This result replicates the change-direction bias observed in Experiment 1 and rules out the possibility of a response bias. It demonstrates that participants not only reported increases in SF but also perceived them more accurately. The SF-increase bias observed in PSSs during Experiment 1 can most likely be attributed to the lower thresholds for detecting SF increases compared to decreases. Furthermore, the mean change detection threshold in the blank condition was 0.80 ± 0.27 |ΔSF octave units| and 0.77 ± 0.18 |ΔSF octave units| in the no blank condition. We found no difference in thresholds between blank and no blank conditions (t(7) = 0.90, p = 0.3966, BF_10_ = 0.47, anecdotal evidence for H0) in the criterion-free change discrimination task (Fig. 4B).

**Fig. 4.**
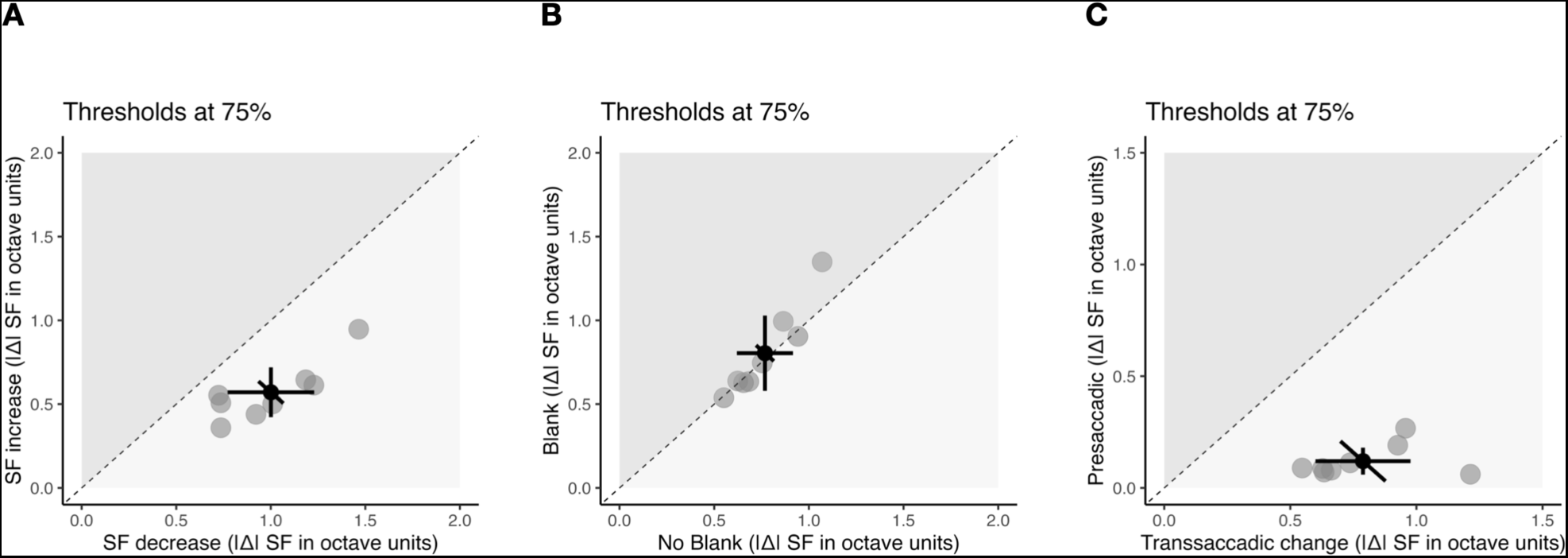
Results of Experiment 2. A) Scatterplot for all detection thresholds compared between SF increase (vertical axis) and SF decrease (horizontal axis) conditions. Data points below the diagonal dashed line indicate lower thresholds (i.e. better performance) in SF increase condition. B) Scatterplot for all detection thresholds compared between no blank (vertical axis) and blank (horizontal axis) conditions. Data points on the diagonal dashed line indicate no difference between the two conditions. C) Scatterplot for all detection thresholds compared between presaccadic (vertical axis) and transsaccadic change (horizontal axis) conditions. Data points below the diagonal dashed line indicate lower thresholds in the presaccadic condition. A - C) Light gray dots represent individual participant data and the dark gray dot indicates the overall mean. The error bars show 95%-confidence intervals for each condition (cardinal bars) or between conditions (oblique bar).

In the presaccadic discrimination task, the mean SF detection threshold was 0.12 ± 0.07 |ΔSF octave units|. We collapsed both change directions in the change discrimination task and conducted a comparison between the thresholds observed in both tasks. Interestingly, the thresholds in the presaccadic discrimination task were significantly lower compared to those in the change discrimination task (t(7) = 8.84, p < 0.0001, BF_10_ = 504.48, extreme evidence for H1) (Fig. 4C). This finding suggests that presaccadically, participants were able to discriminate SF at very low thresholds. Despite that, the precision to detect changes between pre- and postsaccadic stimuli was low, as indicated by the higher thresholds in the first task.

## 4. General Discussion

In this study, we investigated the relationship between visual field differences in appearance before and after a saccade and transsaccadic change discrimination of SF. Our results demonstrate a bias to perceive changes from low to high SF across saccades. A 200 ms postsaccadic blank period improved the precision of the change discrimination but left the bias unaffected. Surprisingly, we could not replicate peripheral overestimation of SF during a saccade; thus, the bias could not be explained by visual field differences. To further explore the distinct effects of presaccadic and postsaccadic information on SF change discrimination performance, we compared thresholds of the presaccadic SF discrimination task (only presaccadic information) and change detection task (presaccadic and postsaccadic information). Lower thresholds in the presaccadic discrimination task could mean that transsaccadic change detection of SF is impaired by overwriting or masking of the presaccadic stimulus by the postsaccadic stimulus. Moreover, there was no observed reduction in detection thresholds in the blank condition when two stimuli were viewed parafoveally. In the following sections, we will discuss each of the findings separately in more detail.

### 4.1 Visual field differences

Our study revealed a uniform SF appearance between peripheral and foveal views. Davis et al. (1987) examined SF appearance at 8 dva along the vertical meridian and found higher apparent SF in the periphery. In our experiment, we examined appearance at 15 dva along the horizontal meridian. Notably, visual performance is generally superior along the horizontal meridian compared to the vertical meridian (for a review see Himmelberg et al., 2023). However, the key distinction from the above studies is that we investigated appearance differences across a saccade, comparing pre- and postsaccadic appearance. This differs from studying peripheral and foveal perception during fixation, as a saccade involves additional mechanisms aimed at aligning pre- and postsaccadic views, thus reducing perceptual disparities (for a review see Stewart et al., 2020). Presaccadic attention shifts that precede saccades, enhance performance at the saccade target (e.g. Deubel & Schneider, 1996). It has been shown that presaccadic attention sharpens visual acuity (Kwak et al., 2023) and shifts SF tuning towards higher SF (Kroell & Rolfs, 2021; Li et al., 2016; Li et al., 2019). The operation of these mechanisms may lead to less significant differences between presaccadic and postsaccadic appearance of SF.

### 4.2 Change direction bias

Although we found no disparities in SF appearance across the visual field, we found a pronounced directional bias in change discrimination. In our second experiment, we ensured that the observed bias was not simply a general preference for one response direction. In addition, we verified that this bias was not due to a limited ability to discriminate SFs in the periphery prior to a saccade. In the periphery, the CSF is shifted, primarily affecting sensitivity to high SFs. This shift may influence the presaccadic perception of high SFs, leading to impaired change perception of SF decrease (SF decrease condition per se implies higher average presaccadic SF compared to the SF increase condition). Nevertheless, our study demonstrated that participants were able to discriminate high SFs presaccadically within a narrow range of 1.68 - 2.38 cpd. The lower thresholds in the presaccadic task compared to the thresholds in the transsaccadic change discrimination task suggest that presaccadic information may be overwritten or masked by postsaccadic information. It should be noted that a direct comparison here is hindered by the different nature of the two tasks. In the presaccadic task, participants discriminated two visually adjacent SFs in the periphery; in the change discrimination task, participants performed discrimination of a memory trace of presaccadic SF information with the postsaccadic SF information.

In the context of object-mediated overwriting (Tas et al., 2012; Tas et al., 2021) our findings indicate that general overwriting occurs because shifts in SF during a saccade without a blank period do not cause significant breaks in object continuity. Furthermore, when there is a shift from high to low SF during a saccade, the effect of a shift on object continuity seems to be even less pronounced and these changes are even less likely to be detected. When the change is accompanied by a blank, the precision of the change discrimination is substantially improved (given there is a single target only). This suggests that the combination of SF change and a blank period disrupts object continuity and protects the presaccadic information from being overwritten. A similar result is predicted by a theory of task-driven visual attention and working memory (TRAM) (Poth & Schneider, 2016; Schneider, 2013), except that, according to TRAM, the establishment of object correspondence across saccades is mediated by attentional weights through prediction. The visual system establishes object correspondence by comparing predicted and actual attentional weights for postsaccadic stimuli. A match increases attention to the postsaccadic information, hindering object feature change detection. A mismatch impairs postsaccadic processing but facilitates change detection due to conflicting attentional weights. Both, object-mediated and attentional weight comparison-mediated overwriting would suggest that change direction discrimination fails due to reduced change detection in general, but they cannot directly explain our change direction bias. Our results show that discrimination of transsaccadic SF change is impaired when SF decreases compared to when it increases. If this perceptual bias does not result from the direct influence of apparent SF or limited ability to discriminate higher SFs prior to saccade, what is its underlying cause? If we look at the bias in terms of segregation and integration of information across saccades, the enhanced change detection for our SF-increase conditions can be interpreted as an increased segregation between two inputs. This may stem from reduced susceptibility of low SF information to postsaccadic masking or decreased masking effectiveness of postsaccadic high SF information. Conversely, diminished change detection in the SF decrease direction suggests that changes were less detectable due to better integration of pre- and postsaccadic information. This may imply increased susceptibility of presaccadic high SF information to postsaccadic masking or enhanced masking capability of postsaccadic low SF information. The latter proposition gains further plausibility when we consider the neural processing of SF information. Visual information is transferred to the visual cortex through two pathways: the magnocellular (M-pathway) for low SF information and the parvocellular (P-pathway) for high SF information. In primates, magnocellular cells display shorter response latencies and higher sensitivity compared to parvocellular cells (Kaplan & Shapley, 1982). Consequently, lower SFs are processed more rapidly than higher SFs (Allen & Freeman, 2006; Hughes et al., 1996). Electrophysiological recordings have verified prolonged peak latencies in evoked potentials activated by higher SF in humans (e.g. Parker & Salzen, 1977). These delays cannot be fully explained by differences in contrast sensitivity alone, since adjusting the perceived contrast of gratings at different spatial frequencies partially relieves these differences, but does not eliminate them (Vassilev & Strashimirov, 1979). Given these differences in processing speed, it is likely that due to changes in SF during the saccade, fast processing is followed by slower processing when low SF information precedes high SF information. Consequently, there would be less overlap between the inputs in terms of processing. Conversely, if high SF information, which is characterized by slower processing, is presented first, followed by low SF information due to the change during a saccade, interference between the two inputs may lead to enhanced masking effects during the transition from high to low SFs. The results of the saccadic latency analysis of the change discrimination task in the first experiment provide evidence for this hypothesis (see Supplementary Material). We found that saccadic latencies increased with increasing presaccadic SF. This observation is also consistent with the findings of Ludwig et al. (2004), who reported longer saccadic latencies associated with high SF Gabors.

The asymmetry between low and high SF information might be related to feedback-processing of peripheral object information in foveal retinotopic cortex (for reviews see Oletto et al., 2023; Stewart et al., 2020): Information about peripheral objects is present in foveal retinotopic cortex (Williams et al., 2008) and the recognition of peripheral objects can be disturbed by TMS over foveal cortex (Chambers et al., 2013) or by delayed presentation of incongruent stimuli in the fovea (e.g. Fan et al., 2016; Weldon et al., 2016). Interestingly, foveal stimuli seem to be more effective in interrupting peripheral recognition if they contain low than high SF information (Goktepe & Schütz, 2024). Given that peripheral processing is focused on lower SF compared to foveal processing (e.g. Chung et al., 2002; Rovamo et al., 1978) the efficiency of a foveal stimulus masking peripheral information depends more on the properties of the peripheral information than on the sensitivity of foveal processing. Hence, postsaccadic, low SF information might be more efficient in interrupting foveal feedback-processing of pressaccadic, high SF information, which would be consistent with the observed change-direction bias.

### 4.3 Blanking

Consistent with previous studies (e.g. Deubel et al., 1996; Goktepe & Schütz, 2023; Grzeczkowski, Deubel, & Szinte, 2020; Grzeczkowski, van Leeuwen, et al., 2020; Hübner & Schütz, 2021; Poth & Schneider, 2016; Tas et al., 2021; Weiss et al., 2015) we observed greater precision in the blanking condition in the change discrimination task of our first experiment. However, the change direction bias was preserved in both blank and no blank conditions. Postsaccadic blanking is known to eliminate or reduce postsaccadic masking effects (e.g. Tas et al., 2021) but here the low-SF postsaccadic stimuli remained to be an effective mask. Interestingly, we did not observe an improvement in detection thresholds in the blank condition of the change discrimination task in the second experiment. One plausible explanation for this outcome is that owing to the nature of the task, participants could not fixate the postsaccadic stimulus foveally; rather, they did so parafoveally (at 2.5 dva), as they were instructed to fixate a cross between the two Gabor gratings. This was different in the change detection task of the first experiment, where observers fixated directly on the Gabors after executing a saccade. Blanking may be less effective when applied to stimuli located in the peripheral visual field, potentially proving most efficacious for saccade targets (but see Deubel et al., 2010). It is also important to consider that this task involves making a saccade between two Gabors, of which only one changes. The unchanging Gabor and a fixation stimulus between the Gabors, which remains visible in the blank condition can act as a reference and potentially influence the perception of the changing target (Deubel, 2004; Deubel et al., 2010).

## 5. Conclusion

We conclude that the peripheral and foveal appearance of a target’s spatial frequency does not differ when a saccade is performed. Nevertheless, the perception of SF changes is persistently biased towards perceiving SF increases across saccades. This bias can even outlast stimulus blanking. Good presaccadic discrimination performance without a postsaccadic stimulus might suggest masking or overwriting by the postsaccadic input to be at play here. Stronger masking/ overwriting seems to occur when the SF decreases during a saccade, resulting in a bias to perceive an increase in SF across saccades.

## Supporting information

Supplementary pdf contains all supplementary files

## Acknowledgements

We thank all participants of the study. The study was supported by Deutsche Forschungsgemeinschaft (DFG), Project number 290878970-GRK 2271 / Project 5 and by the European Research Council under the European Union’s Horizon 2020 research and innovation programme (grant agreement no. 676786).

## Data availability

Psychophysical data and analysis scripts of the experiments will be published upon acceptance at https://doi.org/10.5281/zenodo.10718855.

