## Supplementary pdf contains all supplementary files for "A bias in transsaccadic perception of spatial frequency changes"

### Supplementary material

*1.1 Latency analysis in the change discrimination tasks*

We additionally analyzed saccade latencies in the change discrimination tasks of both experiments. In the change discrimination task of the first experiment, the average mean saccade latency over all participants and conditions was 229.65 ± 50.90 ms (mean ± standard deviation). In the spatial frequency (SF) increase condition the average mean saccade latency was 225.35 ± 24.4 ms. In the SF decrease condition, the average saccade latency was 234.06 ± 22.67 ms. There was a significant difference between the two conditions ($t(16) = -4.0228$, $p = 0.001$, $BF_{10} = 38.36$, very strong evidence for H1). Zimmermann et al. (2013) observed improved intrasaccadic change detection with longer presaccadic observation times. In our data, longer saccadic latencies in the SF decrease condition do not correspond to improved change discrimination, but rather to reduced performance in SF decrease condition. Thus, the differences in saccade latencies between the two conditions cannot explain the differences in discrimination performance in this case.

We also analyzed saccade latency as a function of presaccadic SF (Fig. S1). Simple linear regression was used to test whether presaccadic SF significantly predicted saccadic latency. The overall regression was statistically significant ($R^2 = 0.82$, $F(1, 9) = 40.61$, $p < 0.001$). Longer latencies with higher presaccadic SFs correspond to similar findings by Ludwig et al. (2004), who reported an increase in saccade latency with increasing SF of the Gabors. These results are also consistent with the processing speed hypothesis that we propose in the Discussion.

In the change discrimination task of the second experiment, the mean saccade latency across participants and conditions was 259.78 ± 53.10 ms. In the SF increase condition, the mean saccade latency was 259.06 ± 23.47 ms. In the SF decrease condition, the mean saccade latency was 260.49 ± 20.82 ms. There was no significant difference between the two conditions ($t(8) = -0.7013$, $p = 0.503$, $BF_{10} = 0.39$, anecdotal evidence for H0). No difference between conditions is expected in this task as participants were exposed to two Gabors with distinct SFs before a saccade.

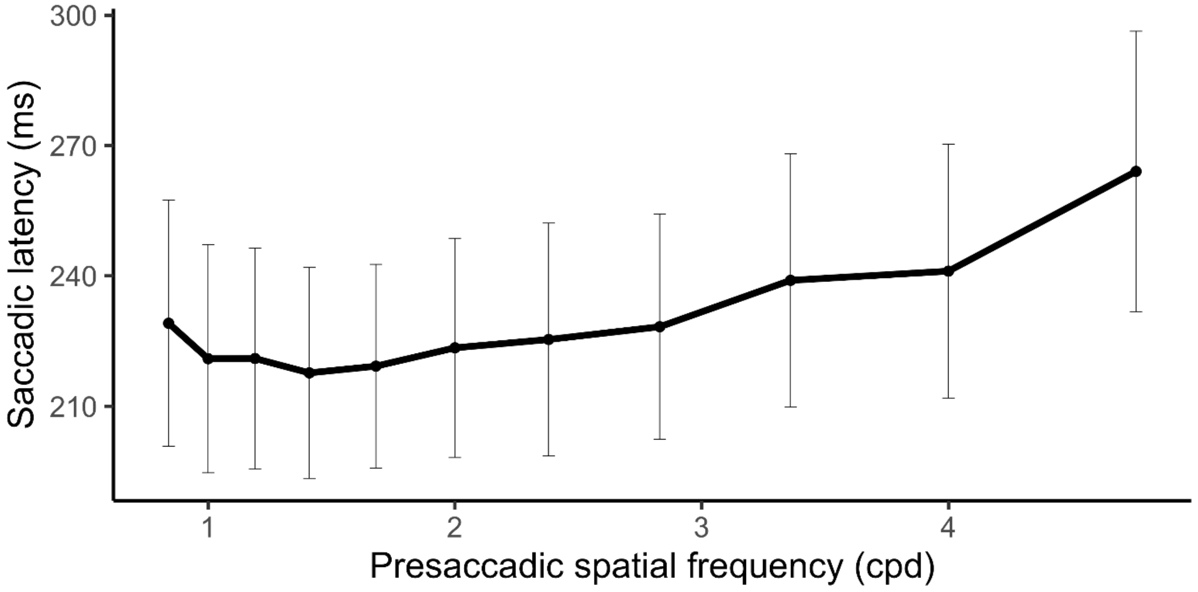


Fig. S1. Saccade latency as a function of presaccadic spatial frequency in the change discrimination task of the first experiment. Rounded presaccadic SFs in cpd on the horizontal axis and mean saccade latencies for each SF on the vertical line. Thin lines indicate the error bars. Participants showed longer saccadic latencies with higher presaccadic SFs.

*1.2 Correlation between PSE and PSS*

Hübner and Schütz (2021) found a correlation between shape appearance differences and change discrimination. Based on these results, we also examined a correlation between PSE difference and PSS, but found no significant correlation between the variables (Fig. S2) (blank condition $r(15) = 0.45$, $R^2 = 0.2$, $p = 0.069$; no blank condition $r(15) = -0.21$, $R^2 = 0.045$, $p = 0.413$).

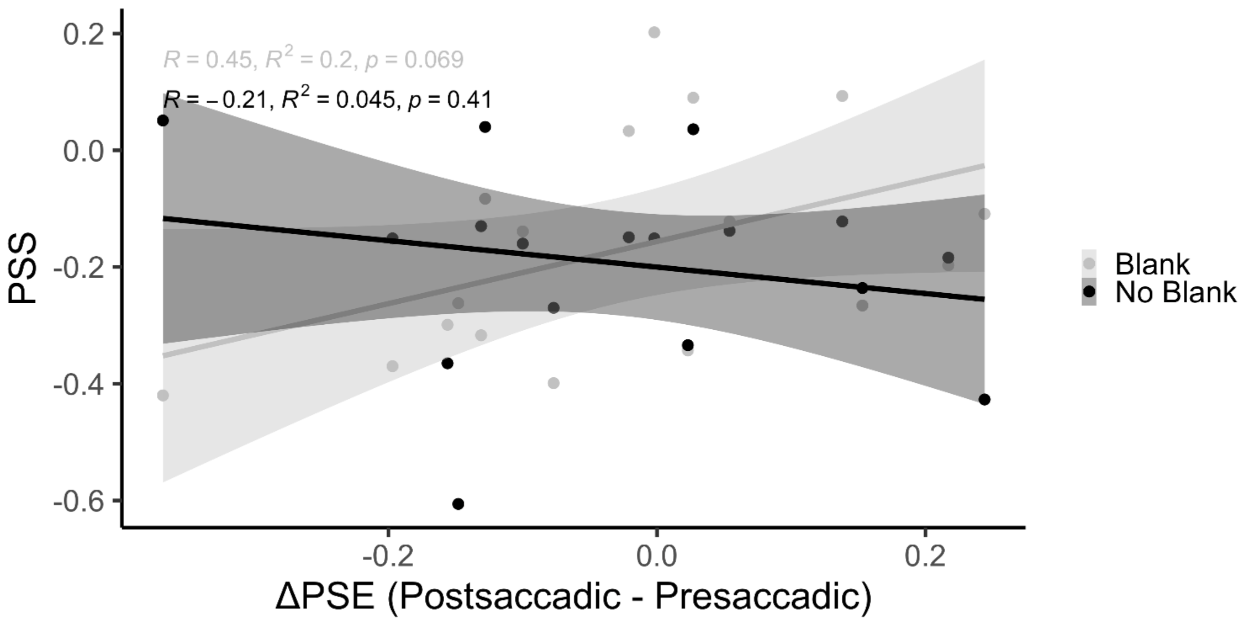


Fig. S2. Differences between presaccadic and postsaccadic PSE (appearance task of the first experiment) on the horizontal axis and change discrimination PSS (change discrimination task of the first experiment) on the vertical axis for the blank (grey) and no blank (black) conditions. Linear regression lines are fitted for both conditions, the shaded areas represent confidence intervals.
